# The scaffolding and activation of NLRP3 on acidic vesicles depends on their biophysical properties and is independent of intermediate filaments

**DOI:** 10.1101/2024.11.18.624139

**Authors:** Guillermo de Paz Linares, Jan Mossemann, Blayne Sayed, Spencer A. Freeman

## Abstract

Inflammasomes are nucleated by receptors that become activated upon cellular stresses including ionic dyshomeostasis. Rather than forming in the cytosol, recent evidence suggests that inflammasomes are nucleated at specific sites in the cell including on cytoskeletal polymers and the membrane surfaces of organelles. The NLRP3 inflammasome, which is formed upon the loss of cytosolic K^+^, had been proposed to form on intermediate filaments as well as on vesicles along the endocytic pathway. To determine the necessary requirement of either mechanism, we used vimentin knockout macrophages which do not have intermediate filaments and compared the formation and function of NLRP3 inflammasomes. We report that vimentin was dispensable for the activation of caspase-1, IL-1β cleavage and release, and inflammatory responses in mice attributed to the inflammasome. Instead, NLRP3 was recruited to PI(3,5)P2, PI(4)P- and LAMP1-positive compartments undergoing osmotic swelling. Swelling of these compartments was dependent on the V-ATPase, the inhibition of which curtailed NLRP3 recruitment and inflammasome activation. Similarly, decreasing the hydrostatic pressure on these vesicles prevented NLRP3 recruitment, IL-1β release and pyroptosis. The results suggest that NLRP3 is activated by biophysical features of acidic organelles in the endocytic pathway.

## Introduction

Inflammasomes are supramolecular organizing centres that assemble in response to diverse threats to cell and tissue homeostasis. Such threats often involve the loss of membrane integrity and/or the breach of the cytosol by microbial-associated molecules (Greaney, Leppla, and Moayeri 2015). The damage that triggers inflammasome activation ultimately leads to cytokine processing and release, which in turn promotes inflammatory cascades and the recruitment of immune cells. Inflammasome assembly is a stepwise process which begins with a transcriptional ‘priming’ step that upregulates expression of its necessary components. This is followed by the sensing of intracellular perturbances via cytosolic pattern recognition receptors and their recruitment of the effector protease, caspase-1. Once concentrated, caspase-1 initiates autolytic cleavage/activation, and subsequent processing of inflammatory mediators including IL-1β and gasdermin D (Brough and Rothwell 2007; He et al. 2015). The NLRP3 inflammasome, which has been extensively studied, is activated in response to insults that ostensibly cause the collapse of the Na^+^:K^+^ gradient between the cytosol and the external medium (Muñoz-Planillo et al. 2013; Perregaux and Gabel 1994; Mangan et al. 2024). Its activation is therefore initiated upon increased permeability of the plasma membrane, which can occur following exposure to diverse bacterial toxins and damage-associated molecular patterns, like extracellular ATP (Franchi and Núñez 2008; Perregaux and Gabel 1994; Laliberte, Eggler, and Gabel 1999). The accumulation of damaging particulates in the endocytic pathway, including alum and monosodium urate crystals (MSU), also triggers NLRP3 inflammasome activation (Hanfen Li et al. 2008; Hornung et al. 2008; Martinon et al. 2006), though mechanisms in this case −and whether these converge on ionic homeostasis− are less clear (Mangan et al. 2024).

Pharmacology aimed at inhibiting NLRP3 has shown enormous potential in mitigating conditions with an underlying chronic inflammatory etiology (Hao Li et al. 2022). Despite the translational relevance of NLRP3 nucleation, recent studies on the location of NLRP3 inflammasome assembly have been seemingly at odds with one another. One report by dos Santos et al., argued that vimentin-based intermediate filaments are required for the assembly of the NLRP3 inflammasome in primary murine macrophages (dos Santos et al. 2015). NLRP3 was found to colocalize and co-sediment with vimentin and the loss of vimentin prevented inflammasome assembly and inflammation. This would seem to be in contrast to other, subsequent reports (Chen and Chen 2018; Zhang et al. 2023; Schmacke et al. 2022) that have found that NLRP3 begins to aggregate on intracellular vesicles enriched for phosphatidylinositol-4-phosphate (PI4P). A stretch of cationic residues found in NLRP3 is thought to facilitate interactions with the negatively charged PI4P. This motif, known as the polybasic region, was essential in mediating the initial assembly of the NLRP3 inflammasome in cell lines (Chen and Chen 2018; Schmacke et al. 2022). Thus, the two models appear to be diametrically opposed, since vimentin is not known to be required for any PI4P-mediated processes.

In this report, we revisit the role of intermediate filaments in the inflammatory response of macrophages, where cytosolic intermediate filaments are solely comprised of vimentin, to NLRP3-activating stimuli. Despite reports of vimentin knockout (*Vim*^-/-^) mice as being protected against a plethora of inflammatory insults (Jiang et al. 2012; dos Santos et al. 2015; Mor-Vaknin et al. 2013; Håversen et al. 2018), we do not find a role for vimentin in the nucleation of the NLRP3 inflammasome nor the subsequent release of inflammatory cytokines from macrophages. Supporting this, we found that *Vim^-/-^* mice were not protected against inflammation in an *in vivo* model of liver ischemia reperfusion injury (IRI), an NLRP3-dependent process (Wu et al. 2022). Therefore, it appears that the protective effects observed in *Vim*^-/-^ mice are attributed to functions independent of NLRP3.

Instead, we find that NLRP3 is recruited to PI4P-, PI(3,5)P_2_, and LAMP1-positive vesicles undergoing osmotic swelling that appear shortly after collapsing K^+^ gradients. NLRP3 recruitment to vesicles (and the resultant IL-1β release) was reversed upon relieving their hydrostatic pressure. Rather than swelling alone being a trigger of assembly, NLRP3 inflammasome activity was dependent on the V-ATPase, implicating K^+^-H^+^ transport on acidic organelles. This suggests that underappreciated biophysical features and ion transport across endomembranes play critical roles in triggering the formation of the NLRP3-inflammasome.

## Results

### ASC specks form independently of vimentin

To test the proposed requirement for vimentin filaments in the nucleation of the NLRP3 inflammasomes, we derived primary macrophages from the bone marrow of *Vim^-/-^*mice. Our bone marrow-derived macrophages (BMDM) did not express vimentin as judged by western blotting (**Fig 1A**) and immunofluorescence (**Fig 1B**). NLRP3-mediated pyroptosis is triggered by the depletion of intracellular [K^+^], including for primary macrophages (Muñoz-Planillo et al. 2013). Experimentally, this is most conveniently achieved with nigericin, a K^+^ ionophore. Alternatively, we also used ATP, which engages purinergic receptors to cause their multimerization and formation of a non-selective pore that conducts K^+^ down its concentration gradient, i.e., out of the cell (Pelegrin and Surprenant 2006). We treated macrophages with nigericin or ATP at standard concentrations and times known to instigate NLRP3 inflammasome formation in a manner dependent on net K^+^-efflux (Perregaux and Gabel 1994; Muñoz-Planillo et al. 2013). Using atomic absorption spectroscopy, we found that the loss of intracellular K^+^ upon the addition of nigericin or ATP was not different between wt and *Vim^-/-^*BMDM, as expected **(Fig. 1C)**. Therefore, any differences in NLRP3 inflammasome activation would not be caused by protection against intracellular [K^+^] depletion in *Vim*^-/-^ BMDM.

**Figure 1.**
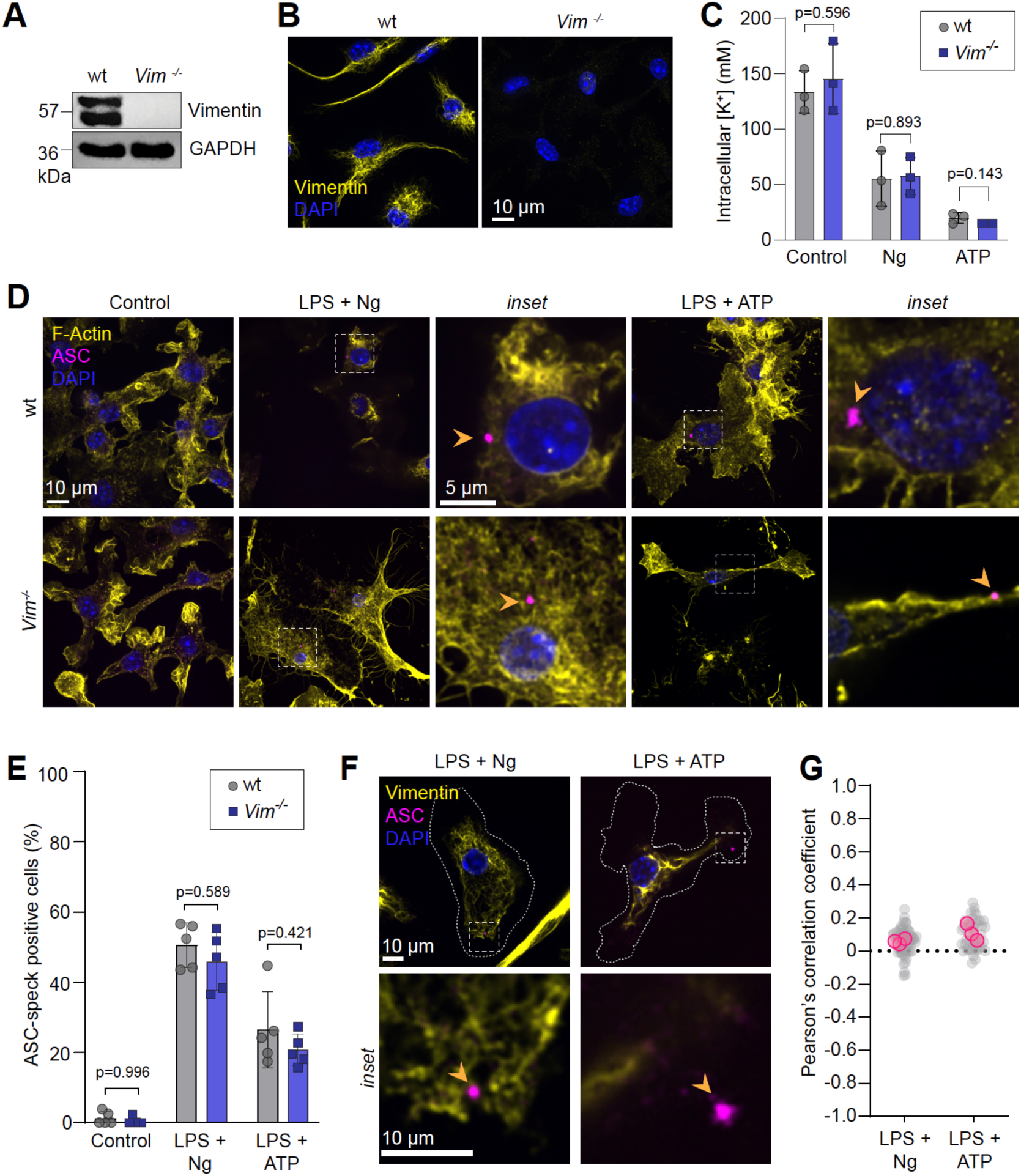
K^+^-dependent ASC aggregation remains unaffected by the absence of vimentin intermediate filaments. Validation of *Vim*^-/-^ BMDM by A) western blot and B) vimentin immunostaining as compared to wt control BMDM. (C) Intracellular K^+^ concentration measurements of BMDM following treatment with nigericin for 15 min or ATP for 30 min. Replicate values ± SD are shown (N=3). (D-G) wt and *Vim*^-/-^ BMDM primed with LPS (1 μg/mL, 4 h) and treated with nigericin (10 μM, 30 min) or ATP (3 mM, 1 h). In (D,E), ASC was immunostained in wt and Vim^-/-^ BMDM, along with F-actin (phalloidin) and DAPI. Representative images shown in D). Data are shown as mean of each replicate ± SEM in E (N=5). In (F,G) cells were immunostained for vimentin and ASC in wt and *Vim*^-/-^ BMDM. Representative images shown in F). (G) Pearson’s correlation coefficient per cell, grey dots and mean, pink dots (N=3, minimum 7 cells per replicate).

Next, we visualized ASC speck formation in lipopolysaccharide (LPS)-primed BMDM, which upregulates the expression of NLRP3 (Bauernfeind et al. 2009), upon K^+^ efflux. ASC is a critical adaptor that enables the recruitment of caspase-1 to NLRP3 and its subsequent activation (Man et al. 2014). We stained endogenous ASC following nigericin or ATP treatment and, surprisingly, found no difference in the incidence of ASC-speck positive BMDM between wt and *Vim*^-/-^ cells **(Fig. 1D,E).** Even more remarkable, when analyzing images using a Pearson’s coefficient, we found there to be no correlation between the localization of vimentin intermediate filaments and ASC specks following NLRP3 inflammasome activation **(Fig. 1F,G)**. This is despite the fact that the intermediate filaments persisted in activated cells. Altogether, these observations would seem to indicate that the assembly of the NLRP3 inflammasome is not dependent on the presence of vimentin intermediate filaments.

### Vimentin is not required for NLRP3-mediated pyroptosis

Once the NLRP3 inflammasome forms, active caspase-1 then cleaves gasdermin D (GSDMD) and pro-IL-1β. Subsequently, the N-terminus of GSDMD inserts into the plasma membrane to form a pore that allows for the passage and release of the processed (mature) IL-1β (Evavold et al. 2018). The involvement of intermediate filaments in GSDMD pore assembly was not assessed in previous studies; conceivably, if vimentin controlled the release of proinflammatory mediators it would explain some results in mice in which KOs are protected from inflammation (Jiang et al. 2012; Håversen et al. 2018; Mor-Vaknin et al. 2013; dos Santos et al. 2015). We treated LPS-primed wt and *Vim*^-/-^ BMDM with nigericin along with propidium iodide (PI) in order to identify potential differences in the rate of plasma membrane permeabilization **(Fig. 2A,B)**. However, we did not see appreciable differences between the vimentin KO macrophages when compared to their wt counterparts.

**Figure 2.**
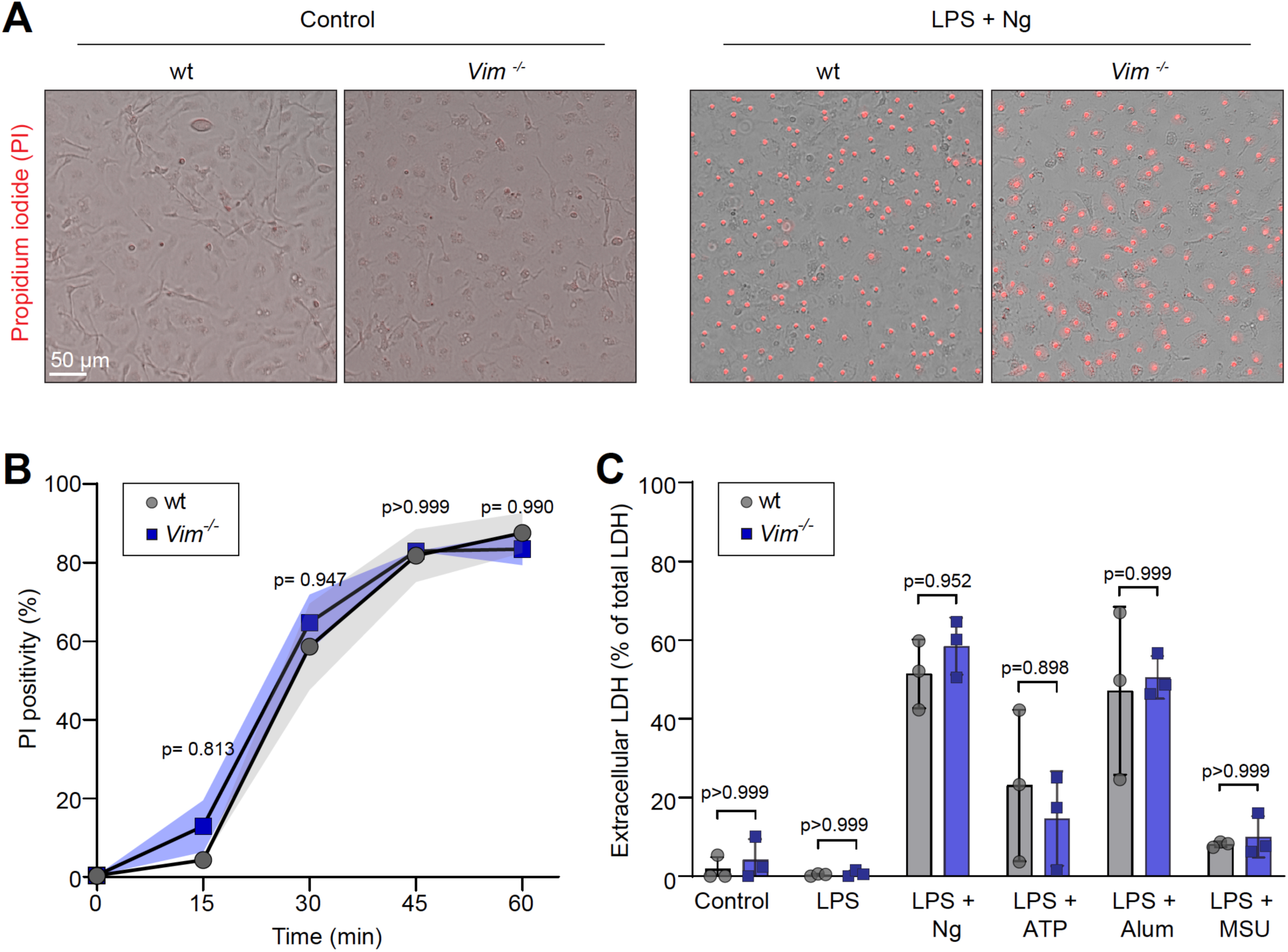
Pyroptotic cell death downstream of NLRP3 inflammasome activation is not dependent on vimentin intermediate filaments. (A,B) LPS-primed (1 μg/mL, 4 h) wt and *Vim*^-/-^ BMDM stained with propidium iodide (1 μg/mL) after nigericin treatment (10 μM) over 1 h. Percent PI positivity is displayed in B) as the mean ± SEM of each timepoint. (N=3). (C) LDH release in wt and *Vim*^-/-^ BMDM primed with LPS (1 μg/mL, 4 h) and treated with nigericin (10 μM) for 30 min, ATP (3 mM) for 1 h, alum (100 µg/ml) for 6 h, or MSU (100 µg/ml) for 16 h. Data represented as replicate mean ± SD (N=3)

Sustained GSDMD pore activity eventually leads to an irreversible form of lytic cell death known as pyroptosis that further promotes IL-1β release into the interstitium (Fink and Cookson 2006; He et al. 2015). Thus, we investigated whether the absence of vimentin in BMDM could prevent or stagnate pyroptosis by measuring the release of lactate dehydrogenase (LDH). In addition to instigating pyroptosis with nigericin or ATP, we also employed alum and MSU crystals to trigger the process. As shown by this assay, nigericin, ATP, alum, or MSU induce lytic cell death in both wt and *Vim*^-/-^ LPS-primed BMDM **(Fig. 2C)**. Moreover, the amount of LDH released from *Vim*^-/-^ BMDM was not significantly different than wt BMDM, irrespective of the NLRP3 inflammasome activator used. Based on our results, we conclude that *Vim*^-/-^ BMDM are not protected against pyroptotic cell death.

### Vimentin is not required for NLRP3-dependent cleavage and release of caspase-1 nor IL-1β

During pyroptosis, caspase-1 activation precedes the cleavage of IL-1β and has been reported to exit the cell as an active protease following cell rupture (Laliberte, Eggler, and Gabel 1999). Indeed, we found cleaved (i.e., active) caspase-1 in the supernatants of LPS-primed BMDM that were treated with either nigericin or ATP **(Fig. 3A)**. Remarkably, we saw that *Vim^-/-^* BMDM were activating and releasing comparable amounts of caspase-1 compared to their wt BMDM counterparts upon pyroptotic stimuli **(Fig. 3A,B)**. Ultimately, IL-1β is a major effector cytokine that results in the propagation of inflammatory responses. We found that BMDM derived from the *Vim^-/-^* mice had no impairment in the upregulation of pro-IL-1β upon LPS stimulation **(Fig. 3C)** and released similar levels of active IL-1β into the supernatant after stimulation with either nigericin or ATP **(Fig 3C)**. Furthermore, we saw no difference in the mature IL-1β concentration found in the extracellular medium of LPS-primed BMDM treated with nigericin or ATP, or upon lysosomal stress (i.e., alum or MSU) **(Fig. 3D)**. Overall, we did not observe any impairment in capase-1 and IL-1β processing and release from *Vim^-/-^*BMDM.

**Figure 3.**
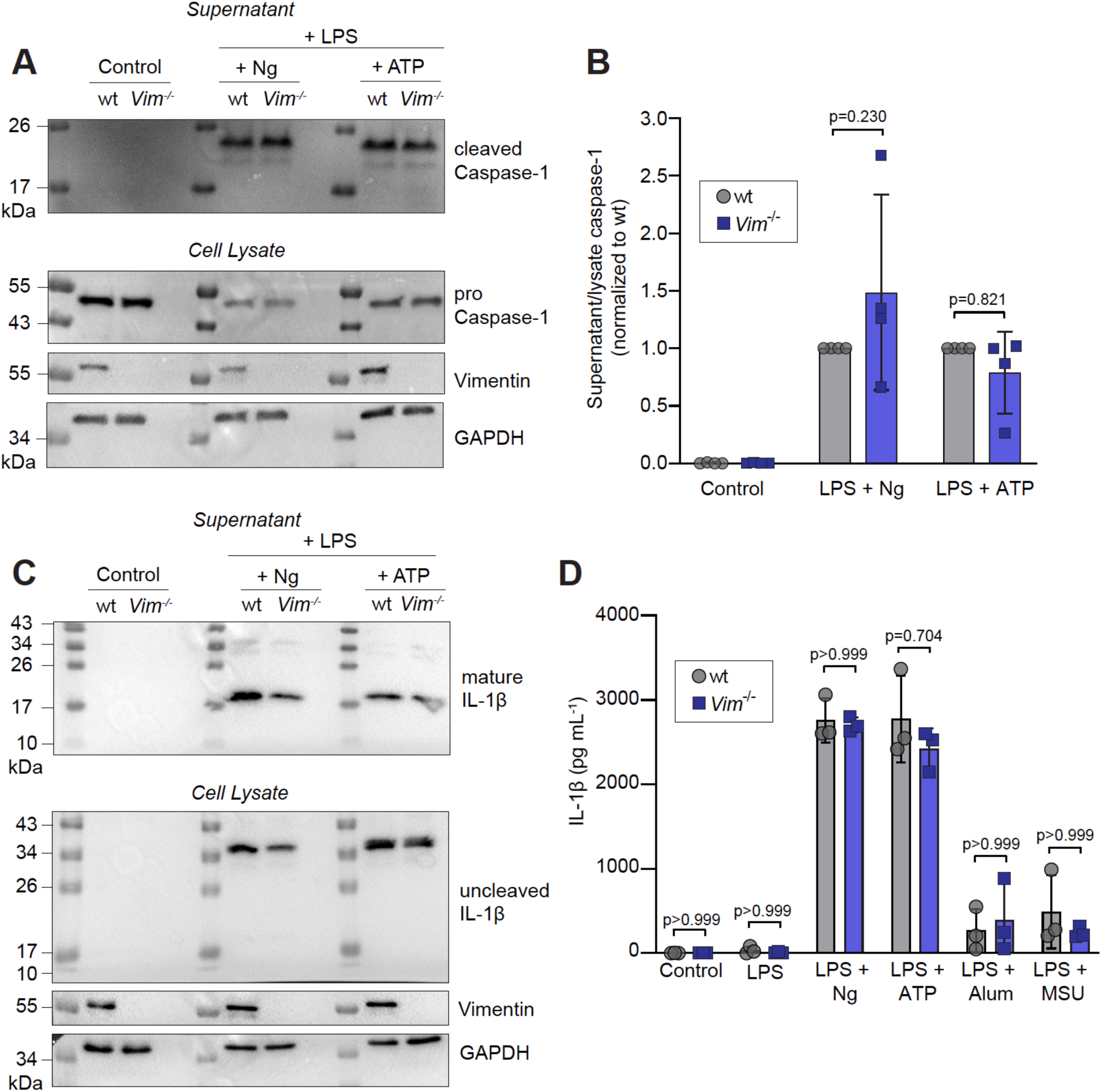
*Vim*^-/-^ BMDM cleave and release caspase-1 and IL-1β following NLRP3 inflammasome activation. (A-C) LPS-primed (1 μg/mL, 4 h) BMDM were treated with nigericin (10 μM, 30 min) or ATP (3 mM, 1 h). Samples were visualized by SDS-PAGE against (A,B) uncleaved and cleaved capase-1, and C) immature and mature IL-1β. In B), quantification of the ratio of cleaved to uncleaved caspase-1 normalized to wt treatments is shown. Individual values ± SD (N=4). (D) Extracellular IL-1β measured in LPS-primed wt and *Vim^-/-^*BMDM treated with nigericin (10 μM) for 30 min, ATP (3 mM) for 1 h, alum (100 µg/ml) for 6 h, or MSU (100 µg/ml) for 16 h. Mature IL-1β concentration was measured by ELISA. Data represented as replicate means ± SD (minimum N=3).

### Vim^-/-^ mice are not protected against hepatic ischemia/reperfusion injury

Despite our *in vitro* observations, there is a plethora of studies reporting that *Vim^-/-^* mice display reduced tissue damage upon numerous inflammatory stimuli (Jiang et al. 2012; Håversen et al. 2018; Mor-Vaknin et al. 2013). In fact, others have reported that *Vim^-/-^*mice are protected against cerebral IRI (Jiang et al. 2012). Thus, we probed whether this protection translated to a hepatic IRI model, which is reportedly mediated, at least in part, by NLRP3 (Jaeschke 2003; Jiménez-Castro et al. 2019; Zhu et al. 2011). Following 1 h of ischemia and 6 h blood reperfusion, we collected samples from mice to look at different aspects of inflammatory injury. First, we did not observe any cohort differences in plasma levels of liver enzymes alanine aminotransferase (ALT) and aspartate aminotransferase (AST) **(Fig. 4A,B)**, indicating that liver damage was not ameliorated in the absence of vimentin. Our histological analyses in which we categorized the percentage of necrotic tissue found in the livers after surgery (*see demarcated yellow areas in* **Fig. 4C**) revealed that, surprisingly, *Vim^-/-^* livers were more necrotic compared to wt **(Fig. 4D)**. Based on these observations, we conclude that the absence of vimentin does not confer a protective effect against the inflammatory damage associated with hepatic IRI.

**Figure 4.**
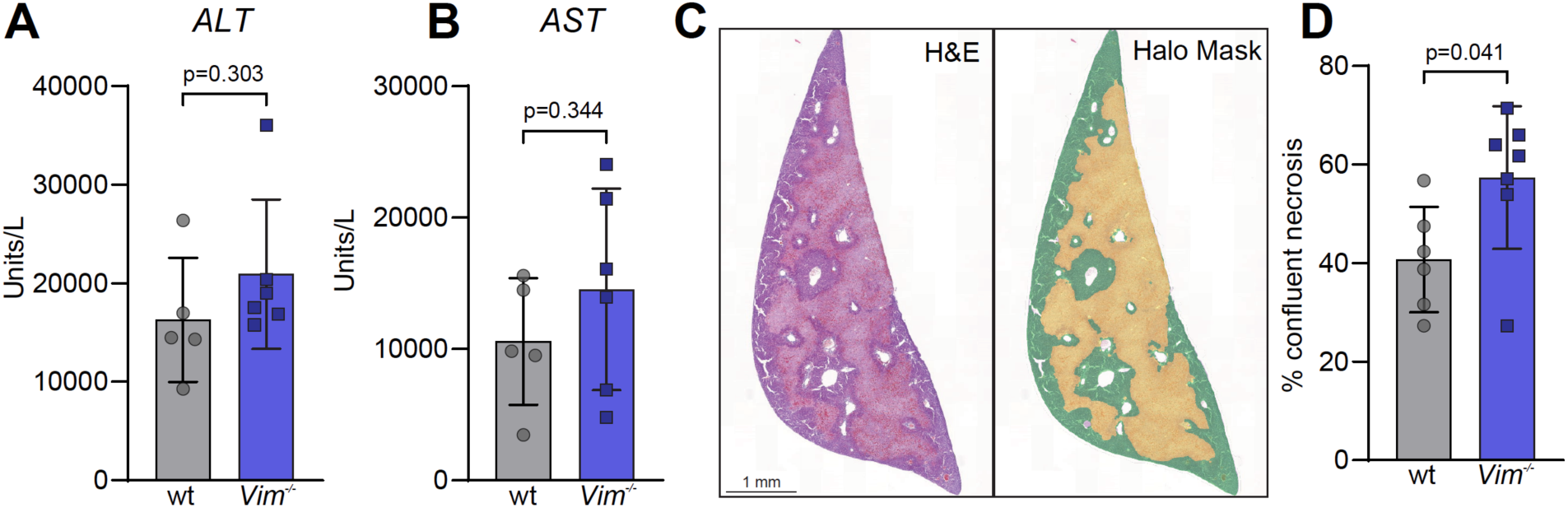
*Vim*^-/-^ mice are not ostensibly protected against hepatic ischemia/reperfusion injury. Warm ischemia/reperfusion injury was modeled using male wt (C57Bl/6) control and *Vim*^-/-^ mice between 6-12 weeks of age. (A,B) Levels of aspartate aminotransferase (AST) and alanine aminotransferase (ALT) in the serum of mice are graphed. (wt N=5, Vim^-/-^ N=6 mice per group.) (C) Hematoxylin and eosin (H&E) staining of liver tissue samples along with Halo DenseNetV2 mask to detect healthy and necrotic areas. (D) Quantification of percent necrosis of liver samples (wt N=6, Vim^-/-^ N=7 mice per group). All data shown as biological replicates ± SD.

### NLRP3 is recruited to PI4P- and PI(3,5)P_2_-containing vesicles

An opposing view to the role of intermediate filaments in scaffolding the NLRP3 inflammasome is that the complex is instead assembled on vesicular membranes requiring electrostatic interactions between NLRP3 and phosphoinositides (Schmacke et al. 2022; Chen and Chen 2018; Zhang et al. 2023). NLRP3 is reported to recognize PI3P, PI4P, and PI(3,5)P_2_ *in vitro*, though only its interaction with PI4P has been demonstrated in cells (Chen and Chen 2018). We therefore expressed an mCherry-NLRP3 fusion protein in HeLa cells, noting that nigericin treatment resulted in its clear enrichment at vesicular structures over the course of 1 hour **(Fig. 5A and Supplemental movie 1)**. Using biosensors for relevant phosphoinositides, we reperformed this experiment and used Statistical Object Distance Analysis (SODA) to determine coincidence of the signals, which provides statistical maps of the assemblies in a population of objects by using morphology and the distance separating objects (Lagache et al. 2018). By this method, we identified objects that were significantly coupled at a radius of 6 pixels from each object centre. We noted no appreciable association between NLRP3 and PI3P (PX-GFP) both at 15 min and at 1 hour **(Fig. 1S)**. Interestingly, by using a recently developed probe for PI(3,5)P_2_, SnxA-GFP, we found clear association between NLRP3 and PI(3,5)P_2_ 15 min post nigericin treatment **(Fig. 1S)**. The association between NLRP3 and PI4P (GFP-P4M) was higher and followed a similar trend to that of PI(3,5)P_2_. As the vesicles increased in size over time, we noted less enrichment of the phosphoinositides on these compartments, which would be consistent with defects in membrane traffic likely occurring well after NLRP3 activation. These data corroborate the finding that NLRP3 assembly is, initially, spatially organized around sites of enriched with PI4P and reveal the possibility that PI(3,5)P_2_ plays a role in NLRP3 activation.

**Figure 5.**
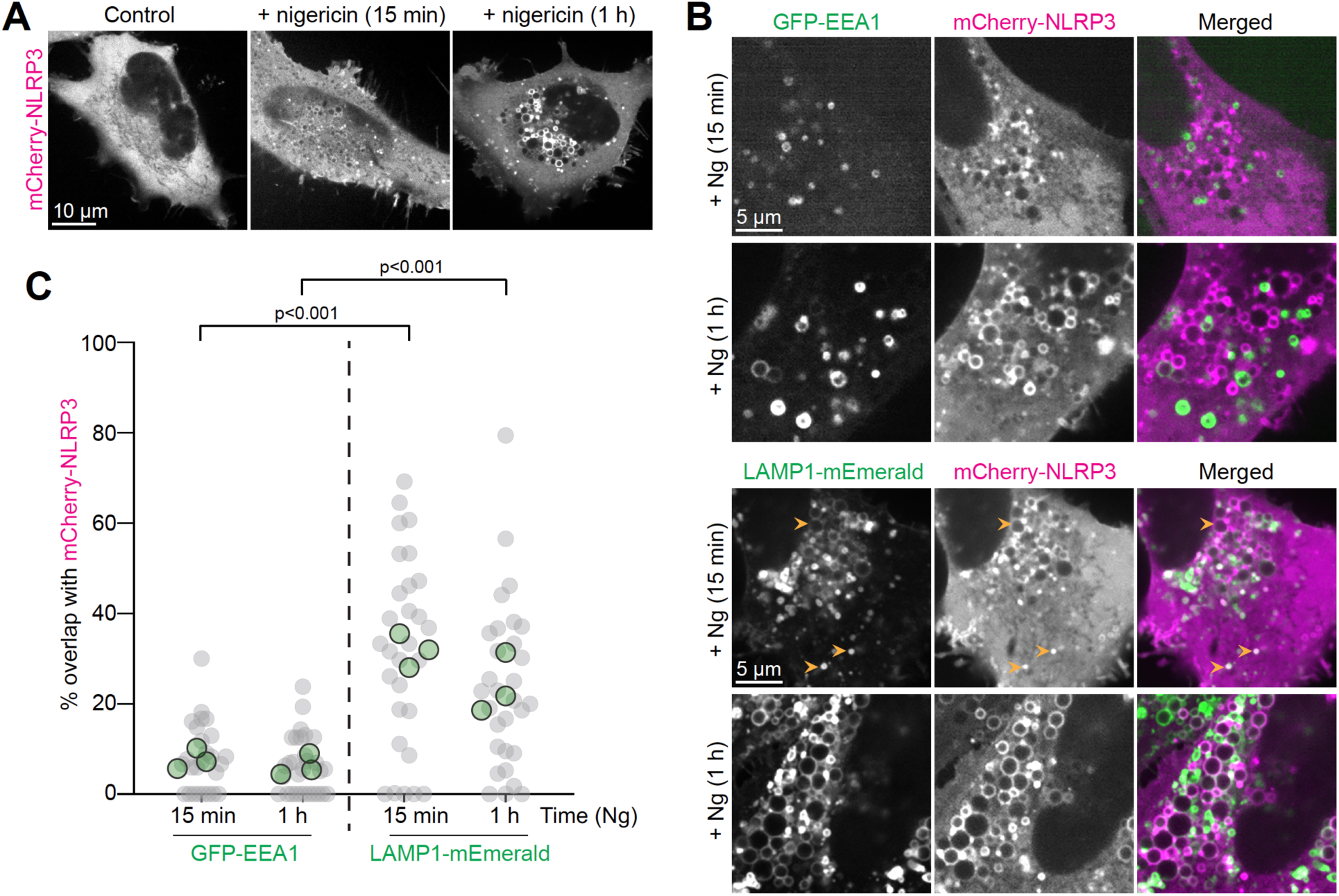
NLRP3 aggregates on LAMP1-positive vesicular compartments. (A) HeLa cells expressing mCherry-NLRP3 before and after treatment with nigericin (10 μM, 15 min and 1h). (B) Co-expression of mCherry-NLRP3 with GFP-EEA1 or LAMP1-mEmerald in nigericin-treated cells after 15 min or 1 h. (C) SODA analysis for object association. Data represented as distribution of percent NLRP3 objects coupled with EEA1- or LAMP1-positive objects. (N=3, with 5-10 cells per technical replicate). Data shown as replicate means and individual technical replicate values.

### Nigericin causes the swelling of LAMP1-positive vesicles that recruit NLRP3

To further characterize the relevant organelle(s) that recruit NLRP3, we co-expressed mCherry-NLRP3 together with early endosomal (GFP-EEA1) and late endosomal (LAMP1-mEmerald) markers. The same SODA analysis was applied which revealed that NLRP3 associates preferentially with LAMP1-positive vesicles following loss of cytosolic potassium **(Fig. 5B,C)**. This association was present at both 15 min and 1 h post nigericin treatment. As with PI(4)P and PI(3,5)P_2_, we noticed that the association of NLRP3 with LAMP1 vesicles once they were swollen tended to be ‘low’ for their expression of LAMP1 while less swollen compartments that associated with NLRP3 were ‘high’ for LAMP1 expression. At later timepoints (i.e., 1 h) NLRP3 associated exclusively with swollen, ‘low’ LAMP1 membrane-bound compartments. This could either reflect the extent to which these vesicles had undergone swelling or the nature of the compartment itself. In contrast, we did not see appreciable NLRP3 aggregation on EEA1-positive compartments at either timepoint. This would suggest that an initiating event in NLRP3 activation occurs at vesicular compartments containing LAMP1 that are beginning to undergo swelling.

### Increases in hydrostatic pressure and the V-ATPase are required for NLRP3 inflammasome assembly and IL-1β release

Swelling of organelles in the endocytic pathway alone would seem unlikely to activate NLRP3; lysosomal storage disorders or osmotic imbalances of endolysosomes do not lead to IL-1β release or pyroptosis (Cai et al. 2024). Supporting this, incubation of cells with a non-digestible, non-transportable disaccharide, sucrose, did not lead to the vesicular recruitment of NLRP3 or pyroptosis, despite leading to the swelling of LAMP1 positive compartments **(Fig. 6A,C).** Vesicular swelling may nevertheless be required for NLRP3 recruitment. To test this, we prevented the gain in hydrostatic pressure incurred upon nigericin treatment by shifting the cells to hypertonic solutions at the time of its addition. The increase in extracellular osmolarity causes water to leave the cell, and organelles, concomitantly (Lang et al. 1998; Hohmann 2002). Under such conditions, we saw the expected inhibition of vesicular swelling. Importantly, preventing the gain in hydrostatic pressure eliminated the recruitment of NLRP3 to vesicular compartments. Instead, NLRP3 remained largely soluble **(Fig. 6B)** and pyroptosis as well as IL-1β release were averted in BMDM **(Fig. 6C,D)**.

**Figure 6.**
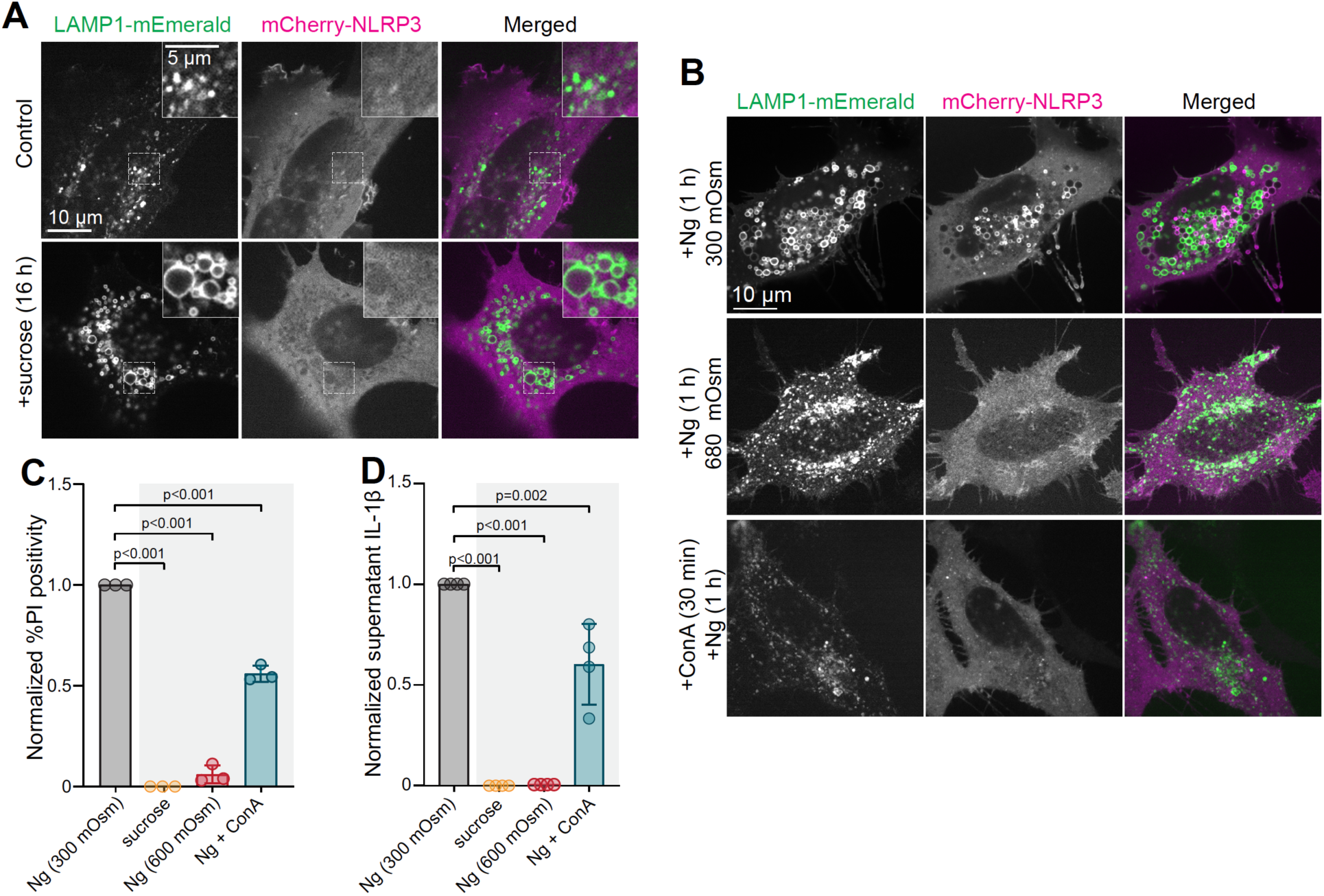
NLRP3 recruitment and activation are dependent on the activity of the V-ATPase. (A) HeLa cells co-expressing mCherry-NLRP3 and LAMP1-mEmerald following treatment with either nigericin (10 μM, 1 h) or sucrose (30 mM, 16 h). (B) HeLa cells co-expressing LAMP1-mEmerald and mCherry-NLRP3 treated with nigericin in either isotonic (300 mOsm) or hypertonic (680 mOsm) medium, or pre-treated with concanamycin A (1 μM, 30 min) prior to nigericin treatment in isotonic medium. (C, D) LPS-primed BMDM treated with nigericin (10 μM, 30 min), sucrose, or concanamycin A (1 μM, 30 min) followed by nigericin. In C), the percent PI positive cells normalized to LPS + nigericin treatment is shown (N=3). In D) the release of IL-1β measured by ELISA and normalized to LPS + nigericin treatment is shown (N=4). All data shown as individual replicate values ± SD.

The swelling caused by nigericin, a membrane-permeable ionophore, may be owed to its direct targeting of endomembranes. Once inserted into the limiting membrane of acidic organelles, nigericin would serve to exchange H^+^ for K^+^, the latter of which may drive swelling. If this is the case, inhibition of the H^+^ gradient (by first inhibiting the V-ATPase) would prevent the accumulation of K^+^. We therefore inhibited the V-ATPase with concanamycin A prior to nigericin addition. This treatment prevented the swelling of LAMP1-positive structures. Critically, this treatment also prevented the recruitment of NLRP3 **(Fig. 6B).** Moreover, concanamycin A pre-treatment led to a reduction in pyroptosis and IL-1β release in LPS-primed BMDM after nigericin addition **(Fig. 6C,D)**. Taken together, we conclude that vesicles endowed with active V-ATPases are the sites of NLRP3 scaffolding and activation upon disruptions to organellar ion transport.

## Discussion

NLRP3 plays a crucial role by acting as a sensor of intracellular dyshomeostasis to then invoke inflammation. K^+^ efflux, indicative of lost plasma membrane integrity or inhibition of the Na^+^/K^+^-ATPase, is a major trigger of NLRP3 inflammasome assembly (Muñoz-Planillo et al. 2013). Solid particles/crystals also activate the NLRP3 inflammasome by causing stress and dysregulation along the endocytic pathway (Hanfen Li et al. 2008; Hari et al. 2014; Groß et al. 2016; Hornung et al. 2008). Whether or not these two, seemingly different stimuli converge mechanistically has not been clear, but this is discussed later on below. Rather than directly sensing differences in alkali cation (Na^+^, K^+^) concentrations in the cytosol, disruption of their normal gradients leads to changes in the cell which are then sensed by NLRP3.

In this study, we demonstrate that vimentin is dispensable for NLRP3-inflammasome assembly and instead find that the V-ATPase and vesicular swelling are required. While many studies implicate vimentin in inflammatory cascades (Mor-Vaknin et al. 2013; Jiang et al. 2012; Eckes et al. 2000; Håversen et al. 2018), this may be owed to impaired immune cell migration and infiltration to inflamed tissues (Barberis et al. 2009; Battaglia et al. 2018; Ivaska et al. 2007; Nieminen et al. 2006). In line with other studies (Zhang et al. 2023; Chen and Chen 2018), our data show that the formation of NLRP3 inflammasomes occurs at vesicular sites enriched in PI4P and, interestingly, PI(3,5)P_2_. Normally these lipids are found in compartments that are distinct from one another, which could be the case here as well since we were not able to image both PIPs together with NLRP3. Alternatively, the immediate impact that nigericin has on membrane traffic could cause the loss of organellar identity in the cells, causing these PIPs to be on the same compartment.

PI4P has recently been shown to mediate other important immune processes pertaining to intracellular danger signals. The cGAS-STING pathway, responsible for sensing cytosolic DNA, was shown to be affected by changes to the intracellular pool of Golgi-PI4P and its associated kinase, PI4KB (Fang et al. 2023; Luteijn et al. 2024). This suggests that the availability of PI4P plays broad and diverse roles in facilitating innate immune responses. Under normal conditions, the majority of PI4P is in the Golgi apparatus and the plasma membrane, as tightly regulated by lipid kinases and phosphatases (Tan and Brill 2014). Nucleation of the inflammasome does not occur at these sites, suggesting that while PI4P may be an essential component of NLRP3 inflammasome activation, there are likely coincidently detected determinants in vesicles that recruit the receptor. The location of PI4P to compartments in which it is not normally found may therefore be an important trigger of inflammation. In contrast to PI4P, little is known about roles for PI(3,5)P_2_ in innate immune sensing and signaling. Lipid-protein overlay assays previously found that NLRP3 can bind PI(3,5)P_2_ (Chen and Chen 2018), though the importance of this interaction remains to be tested.

Others have reported that NLRP3 can bind to vesicles along the endocytic pathway marked with EEA1, Rab5, and/or LAMP1 (Lee et al. 2023; Zhang et al. 2023). We found NLRP3 to preferentially associate with swollen, LAMP1-low vesicles and smaller, LAMP1-high vesicles shortly after nigericin treatment, with this association shifting towards the former over the course of an hour. The nature of these compartments is not yet known. What is clear is that the initial acidification of these vesicles is required for the recruitment of NLRP3. Importantly, dissipation of the H^+^ gradient needs to occur <u>before</u> the addition of nigericin to prevent NLRP3 activation, suggesting that the activity of the V-ATPase is not directly sensed by NLRP3. Nigericin acts as an ionophore for H^+^/K^+^; its insertion into the plasma membrane drives K^+^ loss for H^+^ gain which is an electroneutral process. Acidosis of the cytosol is then relieved by Na^+^/H^+^ exchangers, causing the cell to gain Na^+^. The ionophore also readily inserts into the membranes of organelles. In acidic organelles, the driving forces for ion transport are entirely different from that of the plasma membrane; here, H^+^ will exchange for K^+^ and drive K^+^ into the lumen. In addition to the swelling caused, this also has implications for the electrical potential of the membrane. It is therefore possible that electrical, in addition to biophysical properties of the organelle, are then sensed by NLRP3. While K^+^-efflux is a common trigger of the NLRP3 inflammasome, agents that damage lysosomes and/or disrupt endosomal traffic such as alum, monosodium urate crystals, and silica particles also trigger its assembly (Hari et al. 2014; Groß et al. 2016). Whether or not these damaging agents have similar effects on these electrical/biophysical parameters warrants systematic inquiry.

## Methods and materials

### Animals

*Vim*^-/-^ (129S-*Vim*^tm1Cba^/MesDmarkJ, stock #025692) and wt 129S1 mice (129S1/SvImJ, strain #002448) were obtained from Jackson Labs as previously described (Colucci-Guyon et al. 1994). C57Bl/6 wt mice from Jackson Labs (strain #000664) were used where indicated. All procedures were approved by the Animal Care Committee at the Hospital for Sick Children and were performed in accordance with regulations from the Animals for Research Act of Ontario and the Canadian Council on Animal Care. Mice were kept at the Animal Facility in the Peter Gilgan Centre for Research and Learning.

### Liver ischemia/reperfusion surgery

Cohorts of 8- to 12-week-old Vimentin knockout and wt (C57Bl/6) control mice were used in a 70% segmental warm ischemia-reperfusion model (Selzner et al. 2003). In brief, a sagittal midline laparotomy was made under isoflurane anesthesia, and an atraumatic vascular clamp was placed on the portal vein and the hepatic artery to block blood flow to the left and medial lobes of the liver which account for about 70% of the liver parenchyma. The clamp was removed after 1 h to allow for reperfusion. Following 6 h of reperfusion, mice were euthanized before blood and tissue collection for analysis. Male mice were used to control for known sex differences in IRI responses (W. Li et al. 2018; Ostadal et al. 2019).

### Reagents

Mammalian expression vectors were obtained from the following sources: 2xSidM-GFP as described in (Levin et al. 2017); pCAG mEmerald-LAMP1 was a gift from Franck Polleux (Addgene plasmid # 168513); Human NLRP3 from pEGFP-C2-NLRP3 (gift from Christian Stehlik, Addgene plasmid # 73955) was inserted into an mCherry-C2 mammalian vector by Gibson assembly cloning (NEB, cat #E2611S); pJSK659 (SNX-A-GFP) was a gift from Jason King (Addgene plasmid # 205128); and GFP-EEA1 was a gift from Silvia Corvera (Addgene plasmid # 42307). Primary antibodies used were rabbit anti-cleaved caspase-1 (Asp296) (E2G2I) (Cell Signaling, cat #89332), mouse anti-ASC (BioLegend, cat #676502), rabbit anti-vimentin (Cell Signaling, cat #5741), rabbit anti-pro caspase-1 + p10 +p12 (Abcam, cat. #ab179515) and rabbit anti-GAPDH (Cell Signalling, cat #2118S). Primary antibody validation was done by western blot. Secondary antibodies conjugated with AlexaFluor-405, −488, −555, - 647 and donkey anti-goat HRP secondary antibodies were obtained from Jackson ImmunoResearch. Secondary mouse and rabbit antibodies conjugated with HRP were obtained from Cell Signaling.

Cell culture reagents used include Dulbecco’s modified eagle medium (DMEM, Wisent, cat. #319-007-CL), fetal bovine serum (Atlanta Biologicals, cat. #S12450), Gibco™ DMEM, high glucose, HEPES, no phenol red (FischerScientific, cat. #21063029), HBSS w/out phenol red, calcium and magnesium, supplemented with FBS (Wisent, cat. #311-512 CL), 100x antibiotic-antimycotic solution (Wisent, cat. #450-115-CL), FuGENE 6 Transfection reagent (Promega, cat. #E2691), phosphate buffered saline (Wisent, Sigma-Aldrich, cat#311-010-CL), nigericin sodium salt (Cayman chemicals, cat. #11437), adenosine triphosphate (pH=7.4, cat. #A2383-5G), propidium iodide (Invitrogen, cat. #P1304MP), alum hydroxide (Invivogen, cat. #21645-51-2), monosodium urate crystals (Invivogen, cat. #1198-77-2), lipopolysaccharide (Sigma-Aldrich, cat. # L9764-5MG), concanamycin A (Cayman, cat. # 11050), Prolong Gold Mounting medium with DAPI (Invitrogen, cat #P36935). .

### Cell culture

Bone marrow derived macrophages from the long bones of 6-12-wk-old wt and *Vim*^-/-^ mice was harvested by centrifugation and grown in DMEM + 10% FBS + 10% L929 conditioned medium containing macrophage colony stimulating factor (MCSF) + 1% penicillin-streptomycin at 37°C and under 5% CO_2_. Cells were used 5-7 days after isolation. HeLa cells were grown and passaged in DMEM + 5% FBS + 1% penicillin-streptomycin at 37°C and 5% CO_2_.

### Transfections

HeLa cells were seeded on top of coverslips in 12-well plates and left to attach overnight at 37°C and 5% CO_2_. FuGENE6 Transfection reagent was added to room temperature serum-free DMEM for 5 min, and then plasmid DNA was added at a 1:3 ratio of DNA (μg) to FuGENE6 (μL). Mixture was incubated at room temperature for 15 min and added to each well. Media was changed 4 h post-transfection. Cells were transfected at ∼60% confluency and incubated for at least 18 h at 37°C and under 5% CO_2_ after transfection.

### Intracellular [K^+^] measurements

BMDM were grown to 90% confluency in 2 separate 6-well plates with DMEM supplemented with 10% FBS and 10% MCSF. Following nigericin or ATP stimulation, BMDM from one plate were washed 3x in 150 mM LiCl (pH=7.4) and lysed in 5 mM HNO_3_. Cell lysates were incubated at 4°C overnight and centrifuged at 12,000 g for 15 min to remove any debris.

Samples and standards of known [K^+^] were analyzed using a PinAAcle 900T Atomic Absorption Spectrometer (PerkinElmer, Waltham, MA, USA) to assess the total amount of K^+^ ions. In parallel, BMDM from the second plate were washed as previously described and lifted in isotonic solution. Cell concentration and volumes were determined electronically using a Z2 Coulter particle count and size analyzer (Beckman Coulter) to calculate total cell number and volume for each sample. Intracellular concentrations of K+ were calculated by dividing the total ion content by the total cell volume of each well.

### Immunostaining

BMDM were grown to 70-80% confluency in 12-well plates with DMEM supplemented with 10% FBS and 10% MCSF. Then, BMDM were primed with LPS (1 μg/mL, 4 h) and subsequently treated with nigericin (10 μM, 30 min) or ATP (3 mM, 1 h). Cells were then fixed with 4% PFA for 15 min, permeabilized in 0.1% triton for 5 min and blocked in 5% BSA overnight. Primary and secondary antibodies were diluted to 1:100 and 1:500 in 2.5% BSA in PBS, respectively, and added to coverslips at room temperature for 1 h. Coverslips were washed 3 times with PBS following incubation. Slides were mounted using mounting media containing DAPI.

### PI assay

BMDM were grown to 70-80% confluency in 12 well plates with DMEM supplemented with 10% FBS and 10% MCSF. Cells were primed with LPS (1 μg/mL) for 4 h before simultaneously adding nigericin (10 μM) and propidium iodide (0.3 μg/mL). Pictures of each well were taken in both brightfield and RFP setting of an EVOS microscope (ThermoFischer). At least 3 images of different fields of view were obtained per well per timepoint. Quantification of PI positivity was done in ImageJ using the “Cell counter” feature.

### LDH assay

Cells were grown to 80-90% confluency in 12 well plates with DMEM supplemented with 10% FBS and 10% MCSF. Priming with LPS (1 μg/mL) was done for 4 h, then cells were treated with nigericin (10 μM) for 30 min, ATP (3 mM) for 1 h, alum (100 µg/ml) for 6 h, or MSU (100 µg/ml) for 16 h in 0.1% FBS DMEM. The supernatants of each well were collected and centrifuged at 13,000 g for 5 min, and cells in each well were washed with 1x PBS and lysed in RIPA buffer with protease inhibitor. LDH release was assayed according to manual instructions (CyQUANT LDH Cytotoxicity Assay, Invitrogen, cat #C20301).

### Western blotting

BMDM were grown to 80-90% confluency in 6 well plates with 10% FBS and 10% MCSF. Cells were primed with LPS (1 μg/mL) for 4 h, then treated in reduced serum medium (0.1% FBS) with either nigericin (10 μM) for 30 min or ATP (3 mM) for 1 h. Supernatants were prepared following the TCA precipitation protocol from (Volchuk et al. 2020). Briefly, samples were centrifuged at 13,000 g for 5 min at 4°C to remove any debris and precipitated using 100% TCA. Samples were then washed with ice cold acetone, then heated at 95° C, and finally resuspended in 2x SDS-PAGE sample buffer. For the cell lysates, cells were washed with 1x D-PBS and lysed with RIPA buffer with protease inhibitors. Lysates were centrifuged at 12,000 g for 15 min at 4°C and prepared after determining protein concentration by BCA assay. Lysate proteins were resolved in a 12% SDS-PAGE gel and supernatant proteins using a 15% SDS-PAGE gel. Proteins were transferred to a PVDF membrane and immunoblotted with the appropriate antibodies. Images were analyzed using ImageJ to determine the adjusted mean gray intensity value of each band over the same area.

### IL-1β Enzyme linked immunosorbent assay (ELISA)

BMDM were grown on 12-well plates to 80-90% confluency using DMEM supplemented with 10% FBS and 10% MCSF. BMDM were stimulated and supernatants were collected and centrifuged for 15 min (13,000 rpm, 4°C). Samples were diluted at a ratio of 1:5 of supernatant to the provided dilution buffer. ELISAs were performed according to the manufacturer’s instructions (Mouse IL-1 beta ELISA kit, Abcam, cat. #ab100705). Sample absorbance was read using a Varioskan LUX Plate Reader 1 w/Dual Injectors (Thermofischer Scientific)

### Microscopy

Cells immunostained for ASC and vimentin, as well as live cell imaging was performed by confocal microscopy using a spinning-disk Zeiss Axiovert 200 M microscope (Quorum Technologies Inc.) with an ORCA-Fusion BT cMOS camera, and a five-line laser module with 405-, 443-, 491-, 561-, and 655-nm lines. The microscope operating software was Volocity v6.3 (Perkin Elmer). Images were acquired as Z-stacks with a 63×/1.4 NA for Vimentin/ASC images and 25x/0.8 NA water objective for ASC-speck quantification images. For video acquisition of NLRP3-expressing HeLa cells, a Zeiss Axio Observer microscope with a PCO camera and four-line laser module with 405-, 443-, 491-, and 561-nm lines was used.

For live cell microscopy, coverslips were placed in a chamber with HBSS and imaged on a pre-warmed stage and 63×/1.4 NA oil objective to 37° C. Coverslips used to acquire movies were placed in a chamber along with phenol red-free DMEM and an environmental control chamber set to 37°C and 5% CO_2_. Movies were acquired as single-slice images over the course of one hour.

### Image analysis

Liver tissue was fixed in PFA and stained with hematoxylin and eosin. Slides were digitalized and ischemic score was determined using the DenseNetV2 identifier available in the HALO software. ASC-speck positivity and Pearson’s correlation analysis were done using the Volocity software over the ROI of individual cells. Pearson’s correlation image thresholding was done according to the (Costes et al. 2004) thresholding method to avoid any bias.

SODA analysis of HeLa cells expressing mCherry-NLRP3 and different probes was done using the free image analysis software called Icy (http://icy.bioimageanalysis.org/) and using a modified protocol from (Lagache et al. 2018). Briefly, detection of bright spots on single slice two-channel images was calibrated using the Wavelet Spot Detector. Then, ROIs were made for each individual cell and the statistical probability that spots from both channels were associated with each other by random chance was calculated (as defined by p > 0.05). Probability was found for different radii (*r)* distances from the centre of each spot. Represented in our figure is the coupled puncta of NLRP3 within *r* = 6 pixels (∼516 nm). For each biological replicate, 5-10 cells were analyzed using SODA per probe used.

### Data analysis

All experiments and data were conducted with at least 3 biological replicates. Data is represented as mean ± SEM or individual replicate values ± SD. Statistical significance was determined using two-tailed unpaired Student’s *t* test (significant difference threshold p < 0.05). For multiple comparisons, two-way ANOVA was used with Tukey’s correction for all wt vs *Vim^-/-^*analysis. All other multiple comparisons were done using a one-way ANOVA with Tukey’s correction. All experiments shown were conducted at least 3 independent times (N≥3). All data analysis was done using GraphPad Prism 6 and Excel.

---

**Supplemental Figure 1.**
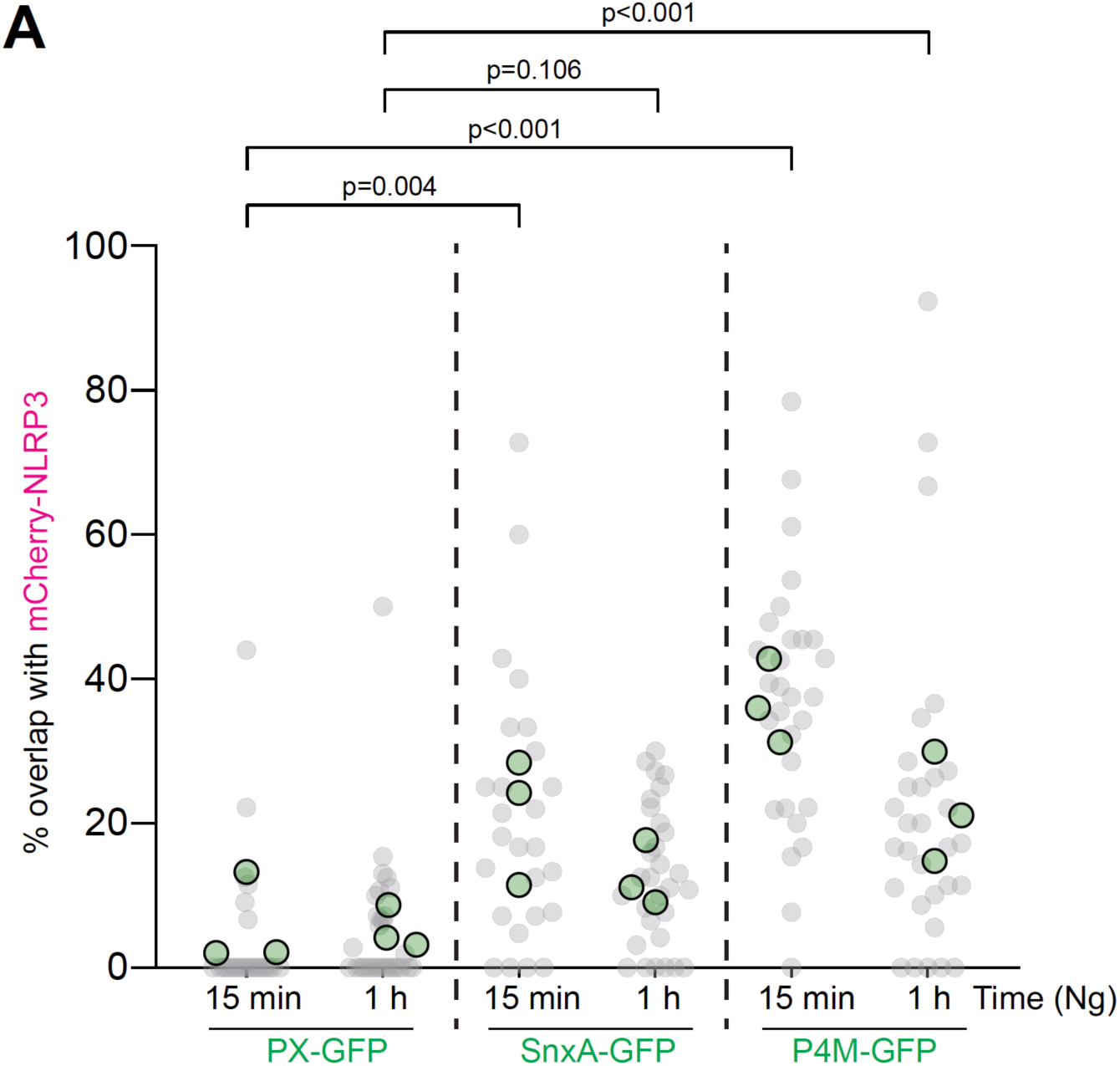
NLRP3 associates with PI4P- and PI(3,5)P_2_-positive vesicles following nigericin treatment. (A) SODA analysis for object association between NLRP3-mCherry and PX-GFP, SnxA-GFP, or GFP-P4M phosphoinositide probes following nigericin treatment (10 μM, 15 min and 1h). Data represented as distribution of percent NLRP3 objects coupled with phosphoinositide probe objects. (N=3, with 5-10 cells per technical replicate). All data shown as replicate means and individual technical replicate values.

